# Synergistic epistasis of the deleterious effects of transposable elements

**DOI:** 10.1101/2021.05.21.444727

**Authors:** Yuh Chwen G. Lee

## Abstract

The replicative nature and generally deleterious effects of transposable elements (TEs) give rise to an outstanding question about how TE copy number is stably contained in host populations. Classic theoretical analyses predict that, when the decline in fitness due to each additional TE insertion is greater than linear, or when there is synergistic epistasis, selection against TEs can result in a stable equilibrium of TE copy number. While several mechanisms are predicted to yield synergistic deleterious effects of TEs, we lack empirical investigations of the presence of such epistatic interactions. Purifying selection with synergistic epistasis generates repulsion linkage between deleterious alleles and, accordingly, an underdispersed distribution for the number of deleterious mutations among individuals. We investigated this population genetic signal in an African *Drosophila melanogaster* population and found evidence for synergistic epistasis among TE insertions, especially those expected to have large fitness impacts. Curiously, even though ectopic recombination has long been predicted to generate nonlinear fitness decline with increased TE copy number, TEs predicted to suffer higher rates of ectopic recombination are not more likely to be underdispersed. On the other hand, underdispersed TE families are more likely to show signatures of deleterious epigenetic effects and stronger ping-pong signals of piRNA amplification, a hypothesized source from which synergism of TE-mediated epigenetic effects arises. Our findings set the stage for investigating the importance of epistatic interactions in the evolutionary dynamics of TEs.

## Introduction

Transposable elements (TEs) are genetic elements that copy themselves and move to new genomic locations (Wells and Feschotte 2020). Their replicative nature and generally harmful impacts on host functions (Langley *et al.* 1988; Montgomery *et al.* 1991; Maksakova *et al.* 2006; Hollister and Gaut 2009; Bellen *et al.* 2011; Rebollo *et al.* 2011; Robberecht *et al.* 2013; Lee 2015) make TEs commonly known as “genomic parasites.” To counteract the selfish replication of TEs, a process that depends on the transcription of TE sequences, various hosts have evolved mechanisms to transcriptionally or post-transcriptionally silence TEs (Yang *et al.* 2017 p.; Ozata *et al.* 2019; Deniz *et al.* 2019). In addition, TEs can be excised from the genome during transposition or through ectopic recombination among repeats within or between TE insertions (Devos *et al.* 2002; Lagemaat *et al.* 2005). While transcriptional and post-transcriptional silencing is expected to limit the selfish increase of TEs in host genomes, mutation accumulation experiments still found an appreciable rate of TE replication (transposition rate, 10^−5^-10^−4^ per copy per genome per generation; (Nuzhdin and Mackay 1995; Maside *et al.* 2000; Pasyukova *et al.* 2004; Díaz-González *et al.* 2011; Adrion *et al.* 2017). Furthermore, this rate of TE increase is at least two orders of magnitude higher than estimated rates of TE excision (Nuzhdin and Mackay 1995; Maside *et al.* 2000; Pasyukova *et al.* 2004; Adrion *et al.* 2017), implying an appreciable net rate of TE increase. At the same time, many eukaryotic genomes only have limited TE abundance (e.g., less than 1% in honeybee (Wells and Feschotte 2020)). Together, these facts pose an outstanding question— how is TE copy number contained in host populations?

Selection against the deleterious fitness effects of TEs has been theoretically proposed as an answer to this puzzle, as it can be a potent evolutionary mechanism counterbalancing the selfish replication of TEs in natural populations (Charlesworth and Charlesworth 1983; Charlesworth and Langley 1989; Lee and Langley 2010; Barrón *et al.* 2014). Empirical investigations have supported the idea that most TE insertions are deleterious and removed from the populations by purifying selection. For example, a dearth of TEs in or near coding sequences is observed across taxa (Kaminker *et al.* 2002; Stuart *et al.* 2016; Laricchia *et al.* 2017). TEs also have frequency spectra that are highly skewed towards rare insertions (Nellåker *et al.* 2012; Cridland *et al.* 2013; Kofler *et al.* 2015; Quadrana *et al.* 2016; Laricchia *et al.* 2017), which is typical for deleterious mutations. Classic theoretical analyses suggest that when natural selection removing TEs cancels out TEs’ selfish increase, TE copy number can reach a balance in host populations (Charlesworth and Charlesworth 1983). It was further predicted that, whether TE copy number is *stably* contained in host populations depends on the mode of epistatic interactions among deleterious TE insertions ((Charlesworth and Charlesworth 1983), reviewed in (Kelleher *et al.* 2020; Choi and Lee 2020)). Specifically, when every additional TE exacerbates host fitness with a larger effect, or synergistic epistasis among the deleterious fitness effects of TEs, it is possible to have an equilibrium TE copy number that is stable even with other forces perturbating TE evolutionary dynamics.

Synergism among the fitness effects of TEs has been predicted to arise through two mechanisms. For one, the illegitimate recombination between nonhomologous TE insertions, or ectopic recombination, generates highly deleterious chromosomal rearrangements (Davis *et al.* 1987; Kupiec and Petes 1988; Montgomery *et al.* 1991; Lim and Simmons 1994; Mieczkowski *et al.* 2006). Empirical evidence suggests that selection against ectopic recombination between TEs is a critical force limiting the selfish increase of TEs in host populations (Langley *et al.* 1988; Montgomery *et al.* 1991; Petrov *et al.* 2003, 2011). Because ectopic recombination happens between two TE insertions, the frequency of the event and the resultant decline in host fitness would naturally depend on the square of TE copy number (Montgomery *et al.* 1987; Langley *et al.* 1988). In other words, each additional TE would incur a higher fitness cost, exhibiting synergistic epistasis. For another, TE-induced changes of local chromatin states are also predicted to give rise to synergistic fitness effects (Lee and Langley 2010; Lee 2015). Small-RNA directed enrichment of repressive epigenetic marks at euchromatic TEs has been identified as a near-universal mechanism to transcriptionally silence TEs in multicellular eukaryotes (Aravin *et al.* 2008; Sienski *et al.* 2012; Le Thomas *et al.* 2013; Marí-Ordóñez *et al.* 2013; McCue *et al.* 2015) (reviewed in (Czech *et al.* 2018; Deniz *et al.* 2019)). Interestingly, these repressive marks could spread to TE-adjacent genic sequences, influencing host functions and, accordingly, fitness (reviewed in (Choi and Lee 2020)). Small RNAs that initiate TE-transcriptional silencing are generated from TE transcripts either directly (e.g., in plants (Xie *et al.* 2004; Kasschau *et al.* 2007)) or indirectly (e.g., via feed-forward “ping-pong cycle” in animals (Gunawardane *et al.* 2007; Brennecke *et al.* 2007; Aravin *et al.* 2007)). Accordingly, the tendency of a TE being targeted and epigenetically silenced by small RNAs, and the consequential deleterious spreading of repressive epigenetic marks, is expected to grow quadratically with increased TE copy number, leading to synergism among the fitness impacts of TEs (Lee and Langley 2010; Choi and Lee 2020). Interestingly, due to the differences in molecular mechanisms, deleterious ectopic recombination and epigenetic effects of TEs have different predictions about which types of TEs are more likely to exhibit synergistic fitness effects.

Although synergism among the harmful impacts of TEs has been long predicted to be an important theoretical requirement for the stable containment of TE copy number, empirical investigations for its presence and extent in natural populations are still lacking (reviewed in (Kelleher *et al.* 2020)). A direct test for the proposed synergistic fitness effects would come from associations between TE copy number and individual fitness. Even though there is an overall negative association between the copy number of a specific TE family and measurements of fitness components (Mackay 1989; Houle and Nuzhdin 2004; Pasyukova *et al.* 2004), inferring the underlying mode of epistatic interactions from these data is challenging. Fitness is multifaceted, and it is hard to identify *a priori* fitness components impacted by the synergistic effects of TEs. The mode of epistatic interactions may also depend on environmental conditions (Peters and Keightley 2000; Kishony and Leibler 2003; Killick *et al.* 2006), further complicating experimental approaches to infer epistatic fitness effects directly. And importantly, subtle effects on fitness (e.g., 1%) are challenging to experimentally measure, but are expected to strongly influence the population dynamics of TEs in nature. An orthogonal approach that does not rely on the direct measurement of individual fitness is therefore needed to investigate the predicted synergistic fitness effects of TEs.

To test the presence of synergistic epistasis among single-nucleotide variants, several methods that do not rely on direct measurements of fitness have been proposed. These methods infer the mode of epistasis from the nonrandom clustering of variants either within species (Sohail *et al.* 2017) or between species (Callahan *et al.* 2011). In particular, (Sohail *et al.* 2017) used the correlation between allele frequencies at different sites, or linkage disequilibrium (LD), to demonstrate the presence of synergistic epistasis among loss-of-function single-nucleotide mutations in human and Drosophila populations. To test the predicted synergism among TE insertions, we applied this population genetic framework to TE presence/absence polymorphism in a *D. melanogaster* Zambian population (Lack *et al.* 2015), which was also the focal Drosophila population in (Sohail *et al.* 2017). This population inhabits the likely ancestral range of the species (Pool *et al.* 2012; Sprengelmeyer *et al.* 2020) and would less likely being influenced by recent demographic history, which could create LD between variants even in the absence of epistatic interactions (Ewens and Spielman 1995; Zavattari *et al.* 2000; Rogers 2014). Importantly, these sequenced *D. melanogaster* strains did not go through intensive inbreeding to establish homozygous lines and were sequenced as haploid embryos (Lack *et al.* 2015). Accordingly, TEs that incur large fitness effects would likely still be represented in the data. By developing a bootstrapping framework, we were able to test for the presence of the predicted synergistic epistasis among TEs and infer from which deleterious mechanisms such synergism likely arises.

## Materials and Methods

### Population genomic data

We used DPGP3 Zambian *D. melanogaster* strains that were sequenced with Illumina paired-end short reads (Lack *et al.* 2015). This dataset includes 197 genomes, and we excluded those that were excluded from (Sohail *et al.* 2017) due to an extreme number of SNPs detected (six genomes), a read length smaller than 100bp (four genomes), or being sequenced in two separate runs (six genomes). An additional eight genomes were removed due to too many missing TE calls (see below). In total, 173 genomes were included in our final analysis. A list of genomes included in the analysis could be found in **Table S1**.

### Identification of TEs

Raw reads of DPGP3 genomes were processed by Trim_galore (“Babraham Bioinformatics - Trim Galore!”) to remove adaptors and low-quality sequences. We used TIDAL (Rahman *et al.* 2015) to identify TE insertions in these DPGP3 genomes with respect to Release 6 reference genome coordinates. All possible TE calls, irrespective of coverage ratio (an index for the confidence of a TE call in TIDAL) and from all genomes, were combined to generate a list of potential TE insertions. We excluded INE-1, a TE family that experienced an ancient burst of activities and whose copies are mostly fixed in *D. melanogaster* (Kapitonov and Jurka 2003; Singh and Petrov 2004). We also excluded TEs on the 4^th^ chromosome, which is nearly entirely heterochromatic (Riddle and Elgin 2018). This yielded 39,084 potential TE insertion sites.

We used previously developed approaches in (Lee and Karpen 2017), which was based on (Cridland *et al.* 2013), to call the presence/absence of all TEs in the list of potential insertion sites in DPGP3 genomes, including the genome in which the TE was identified as an insertion by TIDAL. Briefly, following (Lee and Karpen 2017), we aligned processed reads to Release 6 *D. melanogaster* reference genome using bwa with default parameters (Li and Durbin 2010). Sequences that aligned 500bp around identified TE breakpoints were parsed out using samtools (Li 2011) and assembled into contigs using Phrap (Ewing *et al.* 1998) following parameters in (Cridland *et al.* 2013). The assembled contigs were aligned to TE-masked reference genome using blastn (Camacho *et al.* 2009). A TE is identified as absent if a contig is aligned across the TE insertion site. If no contig spanned over the TE insertion site, contigs were blasted to a database of sequences that include canonical TEs and all TEs annotated in the reference genome (retrieved from Flybase). A TE is called present if there were blast hits to TEs and if a contig aligns to the right or left side of the TE insertion site but does not span across the insertion site. All other scenarios were deemed as missing data (i.e., presence/absence status cannot be determined). We excluded TE insertions that are called present, but the contigs aligned to multiple TE families and thus the family identity of the insertion could not be determined. We used this filtering criterion because an important aspect of our analysis relies on TE family identity (see below). In total, this procedure resulted in 25,998 possible polymorphic (presence/absence) TE insertions.

The TE insertion dataset was further filtered with the following criteria. The strong suppression of recombination in pericentromeric regions is by itself expected to generate LD among variants, independently of synergistic epistasis. Accordingly, we excluded TEs in the heterochromatic regions of the genomes (0.5 Mb inward of the epigenetic euchromatin/heterochromatin boundaries identified in (Riddle *et al.* 2011)). Polymorphic inversions account for a large proportion of population structure (Corbett-Detig and Hartl 2012; Huang *et al.* 2014) and could also create LD among variants. We thus excluded TEs in inversions segregating in the DPGP3 genomes (Lack *et al.* 2015), using inversion breakpoints identified from (Corbett-Detig and Hartl 2012; Huang *et al.* 2014). TE insertions within 1kb to each other, are assigned to the same TE family, and have the same presence/absence calls among all individuals could be two separate TE insertions or one TE insertion that was called twice due to the uncertainty of TE breakpoint identifications. Because we could not distinguish these two possibilities, these 443 TEs were also removed. Following the DPGP3 recommendations, we masked genomic regions with residual heterozygosity, identical by descent, or cosmopolitan admixture (Lack *et al.* 2015). TEs in these regions are considered “missing data.” We then excluded eight genomes whose number of missing TE calls were outliers to other genomes (more than 4,000 missing TE calls, see **Table S1**). We further filter out TE insertions that are called missing data in more than 10% of the genomes or are monomorphic (have the same presence/absence calls among individuals). 11,527 polymorphic TEs passed these filtering. Following (Sohail *et al.* 2017), we further restricted our analysis to rare TEs that are present in equal or fewer than five individuals (11,396 TEs).

### Identification of SNP variants

We used genome assemblies of the same 173 strains (see above) from Drosophila Genome Nexus (Lack *et al.* 2015) (in Release 5 reference genome coordinates). We used Flybase annotation 6.07 (converted to Release 5 coordinates by Liftover (https://genome.ucsc.edu)) to parse out the coding sequence of the longest isoform and then identified synonymous, nonsynonymous, and premature stop codon variants. We excluded genes whose annotation in the reference genome contain putative errors (premature stop codon, lacking canonical stop codon, or having a coding sequence length not multiple of three), following (Sohail *et al.* 2017). Multi-allelic variants (a site with more than two alleles), codons with more than two variants (and thus cannot be assigned as either nonsynonymous and synonymous variants), and SNPs with missing data were excluded from the analysis.

### Estimation and statistical significance of *σ*^2^/*V*_*a*_

For both TEs and SNPs, we restricted the analysis to variants with minor allele counts equal to or smaller than five because TEs/SNPs with allele counts higher than this are unlikely to have deleterious fitness effects. The mutational burden for each individual was estimated as the number of minor alleles of the specific type of variants considered in the genome (Sohail *et al.* 2017). *σ*^2^ is estimated as the variance of mutational burden across genomes. Additive genetic variance (*V*_*a*_) was estimated as Σ_*i*_ 2*p*_*i*_(1 − *p*_*i*_), where *p*_*i*_ is the minor allele frequency of locus/TE insertion *i*.

To evaluate the significance of observed *σ*^2^/*V*_*a*_ of TEs, we used bootstrapped synonymous variants to generate a null distribution for *σ*^2^/*V*_*a*_ for the TE dataset. Specifically, we randomly sampled 1,000 sets of synonymous variants that match the TE dataset in three aspects: (1) the number of variants, (2) minor allele counts, and (3) missing data structure, which controls for the number of missing data of a variant with specific minor allele frequency (MAF) and the total number of missing data per genome. Using the simulated empirical distribution of *σ*^2^/*V*_*a*_, we estimated the one-sided *p-value* for *σ*^2^/*V*_*a*_ of TEs being smaller than the null expectation.

To estimate LD between per pair of TEs, we used PLINK (Purcell *et al.* 2007) to compute the pairwise correlation coefficient (*r*^2^) between all pairs of TEs. For TE insertions *i* and *j,* LD between them (*D*_*i,j*_) is computed as 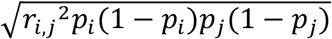, where *p*_*i*_ and *p*_*j*_ are the MAF of TE *i* and *j*. By assuming that TE presence is the derived state, the sign of *D*_*i,j*_ depends on the coupling of TE present alleles, with *D*_*i,j*_ > 0 if TE present alleles are on the same haplotype and *D*_*i,j*_ < 0 for the opposite situation. We then binned pairs of TEs according to their physical distance on the same chromosome or as TEs on different chromosomes (see below), and calculated the mean LD per pair of TEs.

### TE insertions and TE family annotations

To test the predictions that TEs exerting large fitness effects are more likely to show synergistic epistasis, we categorized TEs according to their insertion locations, essentialities of the nearest gene (evolutionary constraints and mutant phenotypes), and local recombination rates. Using Flybase annotation 6.07 and bedtools (Quinlan and Hall 2010), we identified TEs located within exons, UTRs, and introns, and inferred their distance to the nearest gene. To categorize TEs according to evolutionary constraints of nearest genes, we estimated *dN/dS* ratios along the *D. melanogaster* linage using maximum likelihood methods implemented in PAML (v4.9 (Yang 2007)) with alleles from *D. melanogaster, D. simulans* (Hu *et al.* 2013) and *D. yakuba* (Clark *et al.* 2007). Genes with fewer than 100 codons or with *dS* < 0.0001 were treated as missing data. Genes with *dN/dS* estimates were binned into four categories according quartiles of *dN/dS* estimates: [0, 0.0341), [0.0341, 0.0877), [0.0877, 0.1932) and [0.1932, 15.28). To identify genes with essential functions, we used mutant phenotypes identified by either genetic disruptions or RNAi-mediated expression knockdown (downloaded from Flybase 08/22/2018). We focused on three categories related to survival: “lethal,” “semi-lethal,” or “viable.” For genes that have different reported effects on survival, we chose the most severe phenotype. Local recombination rates around TE insertions were interpolated from the estimates of (Comeron *et al.* 2012). We categorized TEs into four bins according to quartiles of local recombination rates (cM/Mbp): [0, 1.344), [1.344, 2.354), [2.354, 3.64), and [3.64, 14.58).

For our analysis that estimated *σ*^2^/*V*_*a*_ of individual TE families, we compared biological attributes of TE families with and without evidence of synergistic epistasis— specifically, their copy number, length, and sequence similarity. TE family copy numbers were estimated from TEs in the reference genome, excluding those in the heterochromatic regions (see above), and from our TE dataset. The mean length of a TE family was estimated by averaging the length of euchromatic copies of the same TE family in the reference genome. To estimate average pairwise sequence difference, we aligned euchromatic TE insertions of the same TE family in the reference genome using MUSCLE (Edgar 2004), calculated the percentage of pairwise difference (excluding gaps), and averaged that over all pairwise comparisons. TEs shorter than 100bp were excluded from the estimation. We also compared TE families for their propensity to be targeted by piRNAs and to exert epigenetic effects. For piRNA-related indexes, we used the estimated amount of sense and anti-sense piRNAs and ping-pong fraction (the proportion of piRNAs generated by the ping-pong cycle) of two wildtype genotypes (*w1118* and *wK*) from (Kelleher and Barbash 2013). For indexes of TE-mediated epigenetic effects of individual TE family (proportion of TEs with the effect, the extent and magnitude of the effect), we used estimates from (Lee and Karpen 2017).

## Results

To investigate the mode of epistatic interactions among TEs using the population genetic framework developed by (Sohail *et al.* 2017), we first identified possible TE insertion positions in the Zambian genomes and then determined the presence/absence status of these TEs in individual genomes. After series of filtering steps to remove TEs with ambiguous family identity or presence/absence status, we identified 11,527 polymorphic TEs in the euchromatic regions of the genome (see Materials and Methods). Consistent with strong selection acting against TE insertions, they had a frequency spectrum that is highly skewed towards rare variants (**Figure 1A**). This frequency spectrum for TE insertions is more skewed than that of SNPs in the same genomes, even those resulting in highly deleterious premature stop codons (Lee and Reinhardt 2012) (**Figure 1A**). Thus, despite few cases of adaptive TEs (e.g., (Daborn *et al.* 2002; Schmidt *et al.* 2010; Hof *et al.* 2016), reviewed in (González and Petrov 2009)), the majority of TE insertions in *Drosophila* appear to be deleterious and are strongly selected against (reviewed in (Charlesworth and Langley 1989; Barrón *et al.* 2014)).

**Figure 1.**
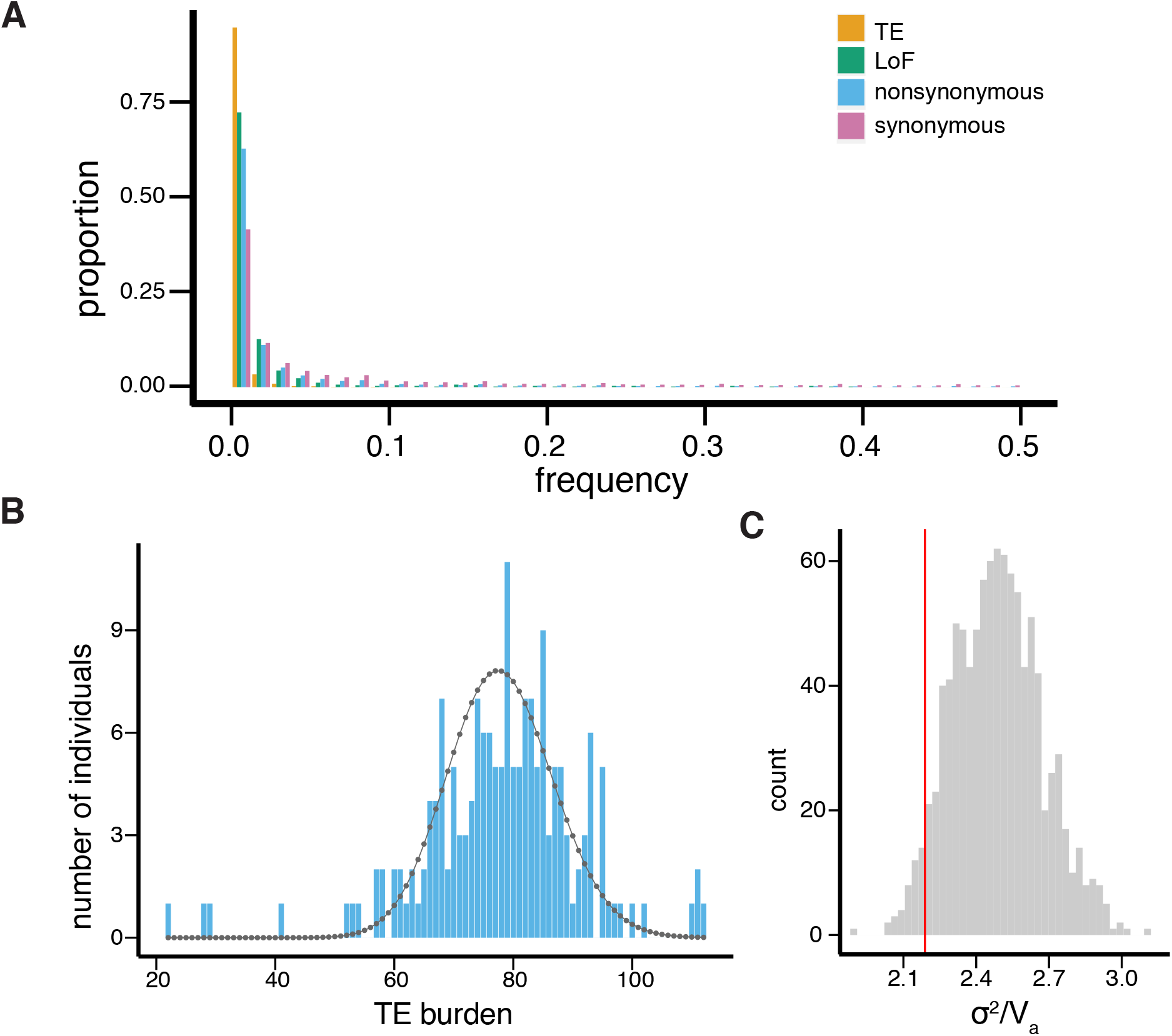
TEs have a skewed frequency spectrum and underdispersed distribution. (A) Frequency spectra of all the TEs that passed filtering and other SNP variants [loss-of-function (LoF), nonsynonymous, and synonymous]. (B) The distribution of TE burden (of TEs with MAF ≤ 5) among 173 genomes. Poisson distribution with identical mean is shown as black dots and line. (C) Resampling distribution of *σ*^2^/*V*_*a*_ for synonymous variants that match number, MAF, and missing data structure as the TE data set is shown in gray bars while the red line shows the observed *σ*^2^/*V*_*a*_ of TEs with MAF ≤ 5.

### Approach for inferring synergistic epistasis among TE insertions

In the absence of epistatic interactions, each mutation decreases individual fitness to the same extent, and selection acts on each mutation independently. Under this circumstance, the variance of mutational burden (*σ*^2^), which could be approximated by the number of deleterious mutations in a genome, would equal the sum of genetic variance across all loci (*V*_*A*_) (Sohail *et al.* 2017). In contrast, with epistasis, there is the interdependency of the fitness effects of mutations, and purifying selection removing them results in LD between alleles (Lewontin and Kojima 1960; Eshel and Feldman 1970; Barton 1995). In particular, selection with synergistic epistasis creates repulsion, or negative, LD among deleterious alleles. The mutational burden will thus have an underdispersed distribution, or smaller variance than would be expected in the absence of epistatic interactions (Charlesworth 1990; Kondrashov 1995).

We estimated “TE burden” as the number of rare TEs in the individual genome (see Materials and Methods). In the absence of other factors that impact the distribution of mutational burden, a reduced variance of TE burden when compared to additive genetic variance (*σ*^2^/*V*_*a*_ < 1) would support synergistic fitness effects of TEs. Yet, even for synonymous variants, which are putatively neutral and should show no epistatic interactions, we found an overdispersed distribution (*σ*^2^/*V*_*a*_ = 7.13). This is similar to previously observed overdispersion of synonymous mutational burden using the same population genomes (Sohail *et al.* 2017), and could result from LD generated by an unknown demographic history of the population or other yet to be identified sources. Further, in our dataset, a large number of TEs are annotated as missing data in at least one genome (99.61%, see Materials and Methods). Missing data is expected to inflate the variance of estimated mutational burden (Sohail *et al.* 2017) and lead to a spurious inference of overdispersion (or *σ*^2^/*V*_*a*_ > 1). Indeed, by randomly masking alleles of synonymous variants, we found a significantly elevated ratio of *σ*^2^/*V*_*a*_ (**Figure S1**). However, excluding TEs with any missing data, which was implemented in previous studies focusing on SNPs (Sohail *et al.* 2017), would reduce the number of polymorphic TEs to only 44 insertions.

Because of these additional factors that could affect the estimated distribution of TE burden, we compared the distribution of TE burden to that of putatively neutral sets of control loci, or synonymous variants. We modified the previously proposed bootstrapping approach (Sohail *et al.* 2017) and randomly sampled sets of synonymous variants that have a matching number of variants, MAF, and missing data structure as the TE dataset. We used these random sets of synonymous variants to generate an empirical null distribution of *σ*^2^/*V*_*a*_ and estimated the *p-value* for the observed *σ*^2^/*V*_*a*_ of TEs being smaller than that of synonymous variants, an approach that assumes that the same factors influence both synonymous variants and TEs.

### TE burden for insertions that likely have large fitness effects has underdispersed distributions

For all euchromatic TE insertions, we found an overall overdispersed distribution (*σ*^2^/*V*_*a*_ = 2.23), and there is an excess of individuals with too large or too small TE burden when compared to Poisson distribution with the same mean of TE burden (**Figure 1B**). Yet, there is also a larger than expected number of individuals with intermediate numbers of TEs (**Figure 1B**). The especially heavy left tail of individuals with too small TE burden could be a result of the abundant missing data for TE insertion (see above). However, despite this overdispersion, TE insertions have significantly lower *σ*^2^/*V*_*a*_ than synonymous controls (*p-value* = 0.043, **Figure 1C**). This observation suggests that TE burden may be underdispersed in the absence of population structure, missing data, and/or other unknown factors inflating the variance. We further partitioned TEs according to TE class and type and estimated *σ*^2^/*V*_*a*_ for each class (Table 1). Among them, the mutational burden of non-LTR (or LINE), retrotransposons that lack long-terminal repeats, has an underdispersed distribution that is significant compared to randomly sampled synonymous variants (*σ*^2^/*V*_*a*_ = 0.98, *p-value* = 0.048; Table 1, **Figure S2**).

**Table 1.** Mutational burden and additive genetic variance of different categories of TEs

| TE type | no. TEs | mean TE burden | $\sigma^2$ | $V_a$ | $\sigma^2/V_a$ | one-sided *p*-value |
| --- | --- | --- | --- | --- | --- | --- |
| All TEs | 11396 | 77.92 | 180.43 | 80.80 | 2.23 | **0.043<sup>a</sup>** |
| **TE class** |  |  |  |  |  |  |
| DNA | 2522 | 17.58 | 25.01 | 18.23 | 1.37 | 0.691 |
| RNA | 8354 | 56.58 | 122.52 | 58.69 | 2.09 | 0.494 |
| **TE types** |  |  |  |  |  |  |
| TIR | 2522 | 17.58 | 25.01 | 18.23 | 1.37 | 0.685 |
| LTR | 6697 | 44.16 | 88.67 | 45.84 | 1.93 | 0.639 |
| nonLTR | 1657 | 12.42 | 12.61 | 12.84 | **0.98<sup>b</sup>** | **0.048** |
| **TE insertion locations** |  |  |  |  |  |  |
| coding exons | 120 | 0.73 | 0.74 | 0.76 | **0.97** | 0.493 |
| UTRs | 5609 | 38.26 | 57.86 | 39.65 | 1.46 | **0.043** |
| exons | 5729 | 38.99 | 66.19 | 44.90 | 1.47 | **0.013** |
| intron | 1498 | 10.49 | 12.24 | 10.85 | 1.13 | 0.381 |
| intergenic | 4169 | 28.44 | 40.97 | 29.54 | 1.39 | 0.172 |
| **dN/dS ratio of nearest gene** |  |  |  |  |  |  |
| 1st quartile: [0, 0.0341) | 2538 | 17.50 | 11.82 | 12.16 | **0.97** | 0.098 |
| 2nd quartile: [0.0341, 0.0877) | 2702 | 18.48 | 13.84 | 12.78 | 1.08 | 0.080 |
| 3rd quartile: [0.0877, 0.1932) | 2840 | 19.35 | 15.86 | 13.35 | 1.19 | 0.058 |
| 4th quartile: [0.1932, ) | 1686 | 11.40 | 8.75 | 7.99 | 1.09 | 0.364 |
| **known pheno.<sup>c</sup> of nearest gene** |  |  |  |  |  |  |
| lethal | 4351 | 9.75 | 40.25 | 30.84 | 1.31 | **0.045** |
| semi-lethal | 827 | 5.64 | 6.23 | 5.83 | 1.07 | 0.487 |
| non-viable (lethal and semi-lethal) | 5178 | 35.38 | 53.88 | 36.66 | 1.47 | 0.116 |
| viable | 4116 | 28.09 | 43.89 | 29.12 | 1.51 | 0.449 |

<sup>a,b</sup> $\sigma^2/V_a < 1$ (<sup>b</sup>) or bootstrapping *p*-value < 0.05 (<sup>a</sup>) are in bold
<sup>c</sup> phenotype

| TE type | no. TEs | mean TE burden | $\sigma^2$ | $V_a$ | $\sigma^2/V_a$ | one-sided *p*-value |
| --- | --- | --- | --- | --- | --- | --- |
| **TEs in exons, dN/dS ratio of nearest gene** |  |  |  |  |  |  |
| dNdS [0, 0.0341) | 1500 | 10.21 | 11.62 | 10.59 | 1.10 | 0.298 |
| dNdS [0.0341, 0.0877) | 1666 | 11.35 | 10.74 | 11.75 | **0.91** | **0.028** |
| dNdS [0.0877, 0.1932) | 1762 | 12.10 | 13.46 | 12.52 | 1.08 | 0.138 |
| dNdS [0.1932, ) | 780 | 5.32 | 6.58 | 5.54 | 1.19 | 0.942 |
| **TEs in exons, known pheno. of nearest gene** |  |  |  |  |  |  |
| lethal | 2855 | 19.36 | 26.35 | 20.08 | 1.31 | 0.412 |
| semi-lethal | 438 | 2.99 | 3.08 | 3.10 | **0.99** | 0.385 |
| non-viable | 3293 | 22.35 | 30.30 | 23.17 | 1.31 | 0.228 |
| viable | 2033 | 13.91 | 20.89 | 14.40 | 1.45 | 0.953 |
| **Intergenic TEs, dN/dS ratio of nearest gene** |  |  |  |  |  |  |
| 1st quartile: [0, 0.0341) | 651 | 4.55 | 4.81 | 4.73 | 1.02 | 0.408 |
| 2nd quartile: [0.0341, 0.0877) | 736 | 5.05 | 5.54 | 5.24 | 1.06 | 0.524 |
| 3rd quartile: [0.0877, 0.1932) | 718 | 4.79 | 4.47 | 4.97 | **0.90** | 0.075 |
| 4th quartile: [0.1932, ) | 802 | 5.38 | 5.58 | 5.58 | 1.00 | 0.248 |
| **Intergenic TEs, known pheno. of nearest gene** |  |  |  |  |  |  |
| lethal | 800 | 5.60 | 5.06 | 5.80 | **0.87** | **0.034** |
| semi-lethal | 226 | 1.43 | 1.55 | 1.49 | 1.04 | 0.676 |
| non-viable (lethal and semi-lethal) | 1026 | 7.03 | 7.46 | 7.29 | 1.02 | 0.257 |
| viable | 1671 | 11.38 | 13.71 | 11.81 | 1.16 | 0.388 |
| **Local recombination rate (cM/Mbp)** |  |  |  |  |  |  |
| 1st quartile: [0, 1.344) | 2838 | 19.65 | 28.75 | 20.33 | 1.41 | 0.659 |
| 2nd quartile: [1.344, 2.354) | 2852 | 19.73 | 23.49 | 20.40 | 1.15 | 0.073 |
| 3rd quartile: [2.354, 3.64) | 2849 | 19.09 | 24.13 | 19.79 | 1.22 | 0.174 |
| 4th quartile: [3.64, ) | 2857 | 19.45 | 30.10 | 20.28 | 1.48 | 0.775 |

Strong purifying selection is expected to weaken the overdispersion of mutational burden generated by population structure (Sohail *et al.* 2017). Signals of underdispersion, if present, should more likely be identified with TEs that exert strong deleterious fitness impacts. We thus categorized TEs according to their potential fitness impacts and examined their distribution separately. By classifying TEs according to their insertion locations, we indeed found that TEs inside coding sequences (*σ*^2^/*V*_*a*_ = 0.97, *p-value* = 0.49), UTRs (*σ*^2^/*V*_*a*_ = 1.46, *p-value* = 0.043) or exons (*σ*^2^/*V*_*a*_ = 1.47, *p-value* = 0.013) likely have underdispersed distributions (**Table 1** and **Figure S3**). This is consistent with the expectation that TEs inserting into coding sequences or UTRs could abolish gene function (Bellen *et al.* 2004, 2011). We also categorized TEs according to the evolutionary constraints of their nearest gene. Genes that have low ratios between nonsynonymous to synonymous substitution rates, or *dN/dS* ratio, are highly conserved and generally expected to have essential functions (Larracuente *et al.* 2008; Waterhouse *et al.* 2011). TEs in or near these genes could potentially result in higher fitness costs. Consistently, we found that mutational burden of TEs whose nearest genes have the lowest quartile of *dN/dS* ratio underdisperses (*σ*^2^/*V*_*a*_ = 0.97, *p-value* = 0.098; **Table 1** and **Figure S4**). Similarly, TEs whose nearest genes have known lethal mutant phenotypes have significantly smaller *σ*^2^/*V*_*a*_ than randomly sampled synonymous variants (*σ*^2^/*V*_*a*_ = 1.31, *p-value* = 0.045; **Table 1** and **Figure S5**). If restricting to insertions within exons, there is a significantly underdispersed distribution for TEs near genes that have the second-lowest quartile of *dN/dS* ratio (*σ*^2^/*V*_*a*_ = 0.89, *p-value* = 0.028; **Table 1** and **Figure S6**). TE burden of exonic TEs in genes with semi-lethal phenotypes also underdisperses (*σ*^2^/*V*_*a*_ = 0.99, *p-value* = 0.385; **Table 1** and **Figure S6**).

In addition to TEs inserting into and disrupting exonic sequences, intergenic TEs could impair host fitness through two mechanisms that were predicted to result in synergistic fitness effects of TEs. The illegitimate recombination between nonhomologous TEs could generate highly deleterious chromosomal rearrangements, irrespective of whether TEs insert inside or outside genes (Davis *et al.* 1987; Kupiec and Petes 1988; Montgomery *et al.* 1991; Lim and Simmons 1994; Mieczkowski *et al.* 2006). Assuming that the rates of ectopic recombination closely follow that of homologous recombination (Lichten *et al.* 1987), TEs in high recombing regions of the genomes should be prone to be involved in ectopic recombination. Yet, we did not find TEs in high recombining regions of the genome having an underdispersed distribution (**Table 1**), which fails to support the prediction of the ectopic recombination model. On the other hand, epigenetically silenced intergenic TEs could result in the spreading of repressive epigenetic marks to adjacent sequences, which disrupts gene functions ((Rebollo *et al.* 2011; Lee 2015), reviewed in (Choi and Lee 2020)). Such an effect is likely to be restricted to intergenic TEs that are near essential genes. Supporting this prediction, there is a significantly underdispersed distribution of intergenic TEs near genes with known lethal mutant phenotypes (*σ*^2^/*V*_*a*_ = 0.87, *p-value* = 0.034, **Table 1** and **Figure S6**). It is worth noting that the deleterious epigenetic effects of intergenic TEs could not be entirely disentangled from TE-mediated genetic disruption of regulatory sequences (Kelleher *et al.* 2020; Choi and Lee 2020). Overall, we found that the distribution of mutational burden for TEs that are expected to have large fitness impacts have underdispersed distributions, especially for TEs inside exons or near essential genes.

### TE burden of many families is underdispersed

Both ectopic recombination and epigenetic effects of TEs depend on sequence homology among TE insertions. Mainly copies of the same TE family recombine, and small RNAs generated from a particular TE family are mostly effective on insertions of the very same TE family. Accordingly, both models predict that the synergistic epistasis would arise among insertions of *the same TE family* (Montgomery *et al.* 1987; Langley *et al.* 1988; Lee and Langley 2010; Lee 2015). We thus estimated *σ*^2^/*V*_*a*_ for individual TE families, restricting to those that have at least 20 insertions (46 of 86, or 53.49%, annotated TE families, **Table S1**). Twenty-two out of 46 TE families (47.83%) have *σ*^2^/*V*_*a*_ < 1 (**Table S2**), and that of *Jockey*, an abundant LINE TE family, is significantly reduced when compared to randomly sampled synonymous variants (*σ*^2^/*V*_*a*_ = 0.81, *p-value* = 0.003, **Table S2** and **Figure S6**). Restricting to TEs that are in or near essential genes found an even higher percentage of TE families with *σ*^2^/*V*_*a*_ < 1 (lowest quartile of *dN/dS* ratio: 64.29% of 28 families; with known lethal phenotype: 58.82% of 34 families; with inviable phenotype: 54.29% of 35 families). Among these families, *Roo*, the most abundant TE family in *D. melanogaster* (Kaminker *et al.* 2002; Kofler *et al.* 2015), and *Jockey* have significantly underdispersed distributions when compared to synonymous controls (**Table S2** and **Figure S6**).

To infer the possible source of synergism among TEs, we compared different attributes of TE families with *σ*^2^/*V*_*a*_ smaller or greater than 1. TE families with *σ*^2^/*V*_*a*_ < 1 should be enriched with those that have synergistic epistasis among copies and thus underdispersed distributions. Factors that may confound *σ*^2^/*V*_*a*_ estimates are unlikely to have differential influence across TE families, making this comparison between groups of TE families less likely biased.

Ectopic recombination and epigenetic effect models share some predictions about which TE families are more likely to exhibit synergistic fitness effects. Both models predict that abundant TE families would elicit higher fitness costs per TE copy (Langley *et al.* 1988; Lee and Langley 2010). Because both mechanisms depend on sequence homology, TEs that are long or have high sequence identity with other copies would represent larger targets for both ectopic recombination and small-RNA mediated epigenetic silencing. Accordingly, it is predicted that TE families that are longer in length or have higher sequence identities among copies should more likely to exert synergistic epistasis (Lee and Langley 2010).

We investigated whether TE families with *σ*^2^/*V*_*a*_ < 1 have larger copy numbers, longer lengths, and higher within-family sequence identities. We used two estimates of euchromatic TE copy numbers: from our data, which is representative of natural populations, and from the reference genome annotation (Kaminker *et al.* 2002; Hoskins *et al.* 2015), which is comprehensive. Because we were unable to assemble internal sequences of TEs with the short-read Illumina data of the focused population, we used annotated euchromatic TEs in the reference genome to estimate the average insertion length and sequence divergence of TEs (see Materials and Methods). We found no significant associations between both estimates of TE copy number and whether a TE family has *σ*^2^/*V*_*a*_ smaller than 1 (*Mann-Whitney U test, p-value* > 0.05 for all comparisons, **Table 2** and **Figure 2A**). Similarly, the sequence divergence between TE families with *σ*^2^/*V*_*a*_ < 1 or > 1 are not significantly different, irrespective of whether we restricted the analysis to insertions in or near essential genes (*Mann-Whitney U test, p-value* > 0.05 for all comparisons, **Table2 and Figure 2B**). On the other hand, the length of TE families with *σ*^2^/*V*_*a*_ < 1 are significantly *shorter* than other TE families when restricting the analysis to TE insertions in or near essential genes (*Mann-Whitney U test, p-value* = 0.0095 (lethal phenotypes) and 0.0075 (inviable phenotypes), **Table 2 and Figure 2C**). The same unexpected pattern held when the analysis used canonical, instead of average, TE length (*Mann-Whitney U test, p-value* = 0.021 (lethal phenotypes), 0.0056 (inviable phenotypes), **Table 2 and Figure 2C**). Overall, we did not find that TE families with *σ*^2^/*V*_*a*_ < 1 are more abundant, longer in length, or with higher sequence identifies than other TE families, defying shared predictions of the ectopic recombination and epigenetic effect models.

**Table 2.**
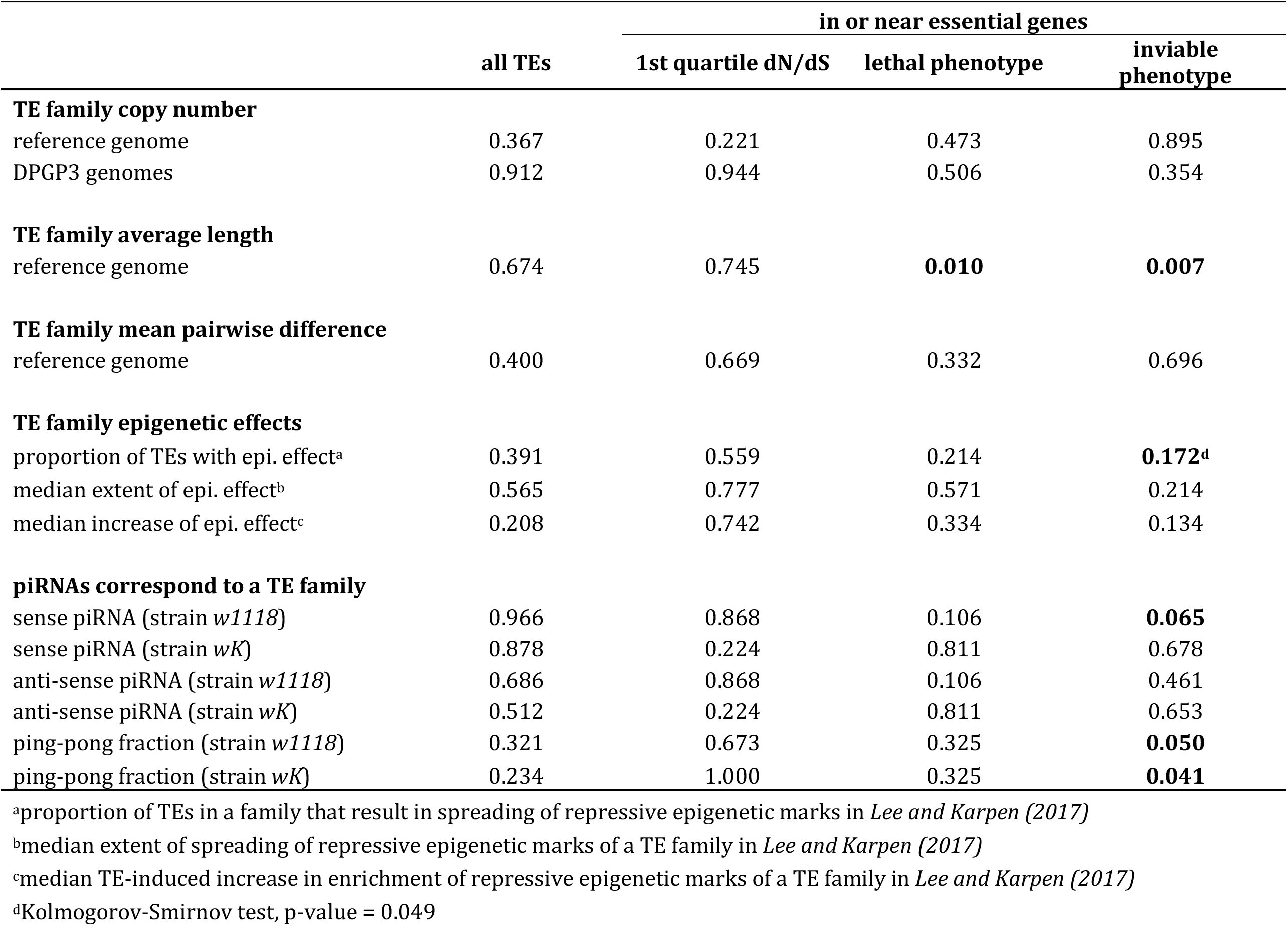
Comparisons of biological properties of TE families with σ 2/Va < 1 or > 1

|  |  | in or near essential genes |  |  |
| --- | --- | --- | --- | --- |
|  | all TEs | 1st quartile dN/dS | lethal phenotype | inviable phenotype |
| **TE family copy number** |  |  |  |  |
| reference genome | 0.367 | 0.221 | 0.473 | 0.895 |
| DPGP3 genomes | 0.912 | 0.944 | 0.506 | 0.354 |
| **TE family average length** |  |  |  |  |
| reference genome | 0.674 | 0.745 | **0.010** | **0.007** |
| **TE family mean pairwise difference** |  |  |  |  |
| reference genome | 0.400 | 0.669 | 0.332 | 0.696 |
| **TE family epigenetic effects** |  |  |  |  |
| proportion of TEs with epi. effect<sup>a</sup> | 0.391 | 0.559 | 0.214 | **0.172<sup>d</sup>** |
| median extent of epi. effect<sup>b</sup> | 0.565 | 0.777 | 0.571 | 0.214 |
| median increase of epi. effect<sup>c</sup> | 0.208 | 0.742 | 0.334 | 0.134 |
| **piRNAs correspond to a TE family** |  |  |  |  |
| sense piRNA (strain *w1118*) | 0.966 | 0.868 | 0.106 | **0.065** |
| sense piRNA (strain *wK*) | 0.878 | 0.224 | 0.811 | 0.678 |
| anti-sense piRNA (strain *w1118*) | 0.686 | 0.868 | 0.106 | 0.461 |
| anti-sense piRNA (strain *wK*) | 0.512 | 0.224 | 0.811 | 0.653 |
| ping-pong fraction (strain *w1118*) | 0.321 | 0.673 | 0.325 | **0.050** |
| ping-pong fraction (strain *wK*) | 0.234 | 1.000 | 0.325 | **0.041** |

<sup>a</sup>proportion of TEs in a family that result in spreading of repressive epigenetic marks in *Lee and Karpen (2017)*
<sup>b</sup>median extent of spreading of repressive epigenetic marks of a TE family in *Lee and Karpen (2017)*
<sup>c</sup>median TE-induced increase in enrichment of repressive epigenetic marks of a TE family in *Lee and Karpen (2017)*
<sup>d</sup>Kolmogorov-Smirnov test, p-value = 0.049

**Figure 2.**
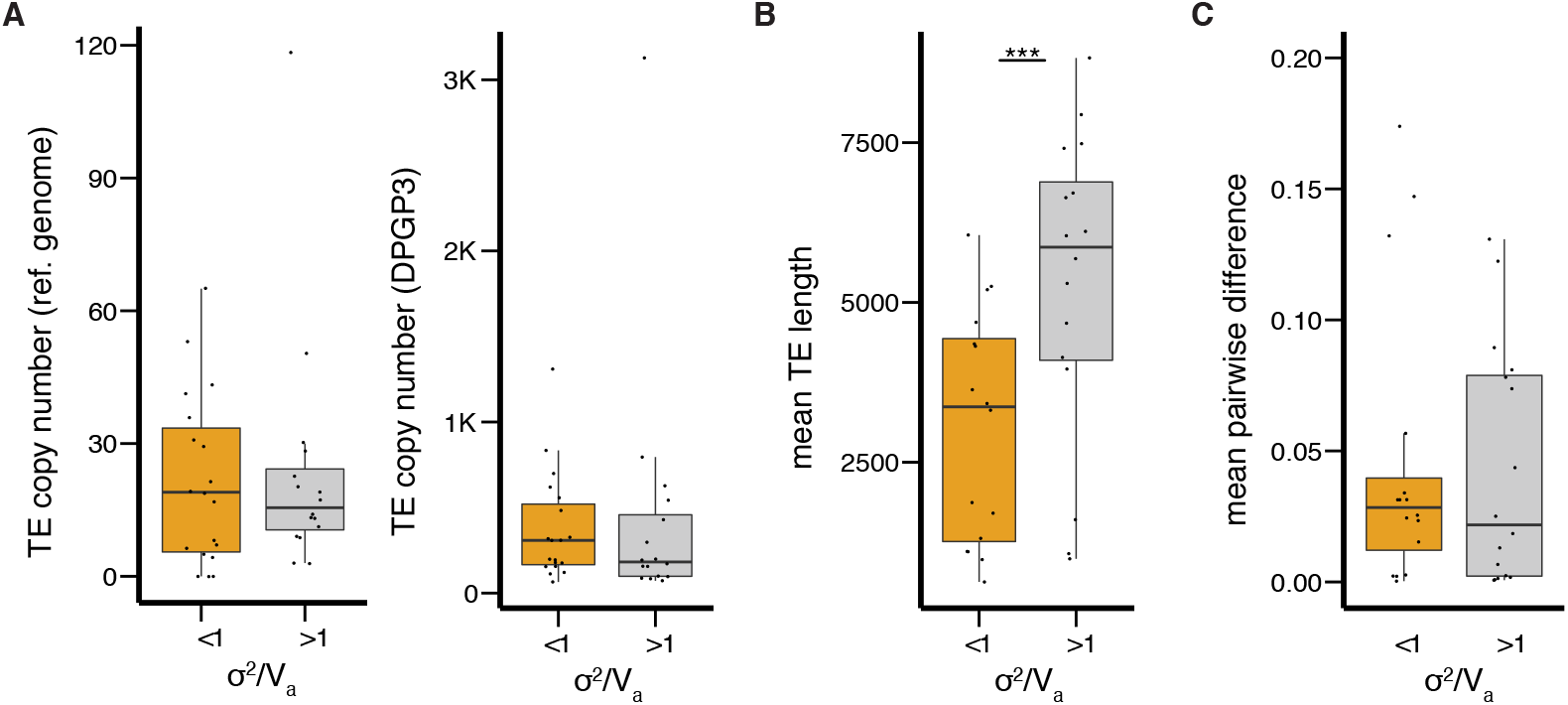
Comparing biological attributes between TE families with *σ*^2^/*V*_*a*_ < 1 or > 1. Boxplots for (A) TE copy number (in the reference genome or in our current dataset), (B) average and canonical TE length, and (C) mean pairwise difference for TE families with and without an underdispersed distribution (*σ*^2^/*V*_*a*_ <1) when restricting the analysis to TEs near/in genes with known inviable mutant phenotypes. Also see Table 2. \*\*\**Mann-Whitney test p-value <* 0.001.

### TE families with stronger epigenetic effects are more likely to have underdispersed distribution

In addition to predictions that are shared with the ectopic recombination model, the epigenetic effect model has several unique predictions about which TE families are prone to exhibit synergistic fitness effects. The propensity to elicit epigenetic effects varies significantly among TE families (reviewed in (Choi and Lee 2020)), and intuitively, TE families that exert stronger such effects are more likely to interact synergistically. The synergism among the deleterious epigenetic effects of TEs in *Drosophila* was predicted to arise through the molecular details for piRNA production (Lee and Langley 2010). While other mechanisms also generate piRNAs ((Malone *et al.* 2009), reviewed in (Czech *et al.* 2018)), “ping-pong cycle” is thought to be responsible for the majorities of the piRNA amplification in flies. In this feed-forward cycle, TE transcripts, which are a source of sense piRNA precursors, and anti-sense piRNA precursors are reciprocally cleaved to generate mature sense and anti-sense piRNAs ((Gunawardane *et al.* 2007; Brennecke *et al.* 2007), reviewed in (Czech and Hannon 2016; Czech *et al.* 2018)). The amount of piRNAs, and accordingly the number of epigenetically silenced TEs and their associated deleterious effects, should grow quadratically or even exponentially with TE copy number (Lee and Langley 2010; Lee 2015). Interestingly, the involvement of ping-pong cycle in the generation of piRNA significantly vary between TE families (Li *et al.* 2009; Malone *et al.* 2009; Kelleher and Barbash 2013). Synergism is expected to have a higher tendency to arise for TE families that are targeted by more piRNAs generated via the ping-pong cycle.

To test these predictions, we used previously estimated indexes for the strength of epigenetic effects for individual TE family: the proportion of TEs resulting in *cis* spreading of repressive marks, median extent of this spreading, and median magnitude of TE-induced increased enrichment of repressive marks (Lee and Karpen 2017). While there is an overall trend that TE families with *σ*^2^/*V*_*a*_ < 1 are more likely to exert epigenetic effects and result in a greater extent of the spreading of silencing marks (**Figure 3A**), only the estimated proportion of TEs with epigenetic effects have a statistically significant shifted distribution (shifted towards large value for families with *σ*^2^/*V*_*a*_ < 1, *Kolmogorov-Smirnov test*, *p-value* = 0.049 when restricting to TEs in/near genes with known inviable mutant phenotypes, **Figure 3A**). We also compared the extent of piRNA targeting between TE families with *σ*^2^/*V*_*a*_ smaller or greater than one (see Materials and Methods). We found that TE families with *σ*^2^/*V*_*a*_ < 1 are generally targeted by more piRNAs (**Figure 3B**). However, none of the comparisons are statistically significant, except for a marginally insignificant higher amount of sense *piRNAs* for TE families with *σ*^2^/*V*_*a*_ < 1 (for TEs in or near genes with known inviable mutant phenotypes, *Mann-Whitney U test, p-value* = 0.065, **Table 2 and Figure 3B**). Interestingly, we found that ping-pong fraction, which estimates the involvement of ping-pong cycle in piRNA generation, is significantly higher for TE families with *σ*^2^/*V*_*a*_ < 1 when focusing on TE insertions near or in genes with known inviable mutant phenotypes (*Mann-Whitney U test, p-value* = 0.050 (*w1118*) and 0.041 (*wK*), **Table 2 and Figure 3C**). Our observations are consistent with the predictions that synergistic epistasis may arise through piRNA amplification from the ping-pong cycle and the resultant deleterious epigenetic effects.

**Figure 3.**
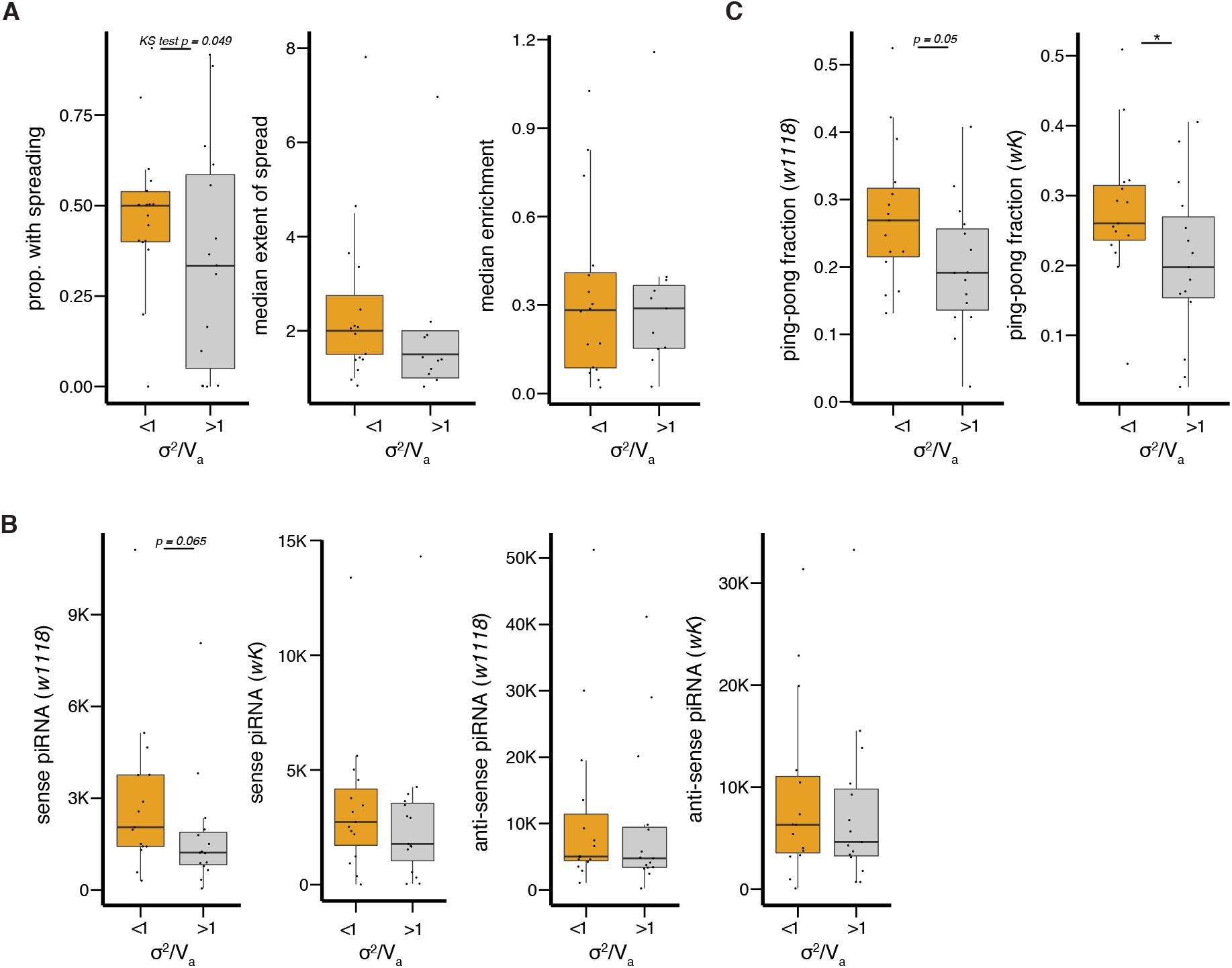
Comparing piRNA-targeting and epigenetic effects between TE families with *σ*^2^/*V*_*a*_ < 1 or > 1. (A) Boxplots for the proportion of TEs showing spreading of repressive marks (left), the median extent of TE-mediated spreading of repressive marks (middle), and TE-mediated increase in the enrichment of repressive marks (right) for TE families with and without underdispersed distributions (*σ*^2^/*V*_*a*_ <1). The distributions of proportion of TEs with spreading effects are significantly different, as indicated by *Kolmogorov-Smirnov test.* (B and C) Boxplots for sense and anti-sense piRNAs (B) and ping-pong fraction (C) corresponding to TE families with *σ*^2^/*V*_*a*_ <1 or > 1 when restricting to TEs that are in/near genes with known inviable mutant phenotypes. The amount of piRNA was estimated as the number of piRNA reads corresponding to a TE family per 1 million TE-corresponding piRNAs in (Kelleher and Barbash 2013). Results of two genotypes (w1118 and wK), both of which are wildtypes for the piRNA pathway genes, are shown. Also, see Table 2. * *Mann-Whitney test, p* < 0.05.

## Discussion

Theoretical analysis has predicted that, to stably contain the selfish increase of TEs, each additional TE insertion needs to impose a higher fitness cost than the last one, leading to purifying selection accelerating the removal of TEs with increased TE copy number (Charlesworth and Charlesworth 1983). This theoretical requirement has been extensively discussed in the context of TE evolutionary dynamics (Charlesworth and Langley 1989; Lee and Langley 2010; Kelleher *et al.* 2020; Choi and Lee 2020), and is predicted to be biologically plausible under several deleterious mechanisms of TEs, including TE-mediated ectopic recombination (Montgomery *et al.* 1987; Langley *et al.* 1988) and the spreading of silencing marks (Lee and Langley 2010; Lee 2015). However, the presence and prevalence of synergistic fitness effects among TEs are yet to be empirically tested.

Purifying selection with synergistic epistasis generates repulsion linkage among variants and, accordingly, an underdispersion of mutational burden when compared to selection without epistatic interactions (Charlesworth 1990; Kondrashov 1995; Sohail *et al.* 2017). Using approaches that leverage this population genetic signal (Sohail *et al.* 2017), we investigated the predicted synergistic epistasis among potentially deleterious TE insertions in the likely ancestral population of *D. melanogaster*. To account for the impacts of known (demography and missing data) and unknown factors that overdisperse TE burden, we compared the distribution of TE burden to that of bootstrapped synonymous variants to identify TEs that are likely underdispersed, an approach that has been implemented in previous studies investigating LD among variants (Sohail *et al.* 2017; Garcia and Lohmueller 2020) (but see (Sandler *et al.* 2020) for potential caveats). An important assumption of this approach is that overdispersing factors would influence the mutational burden of TEs and synonymous variants similarly. Factors that only overdisperse TE burden (e.g., variation in *trans* factors that influences TE silencing between genomes (Lee and Karpen 2017)) could not be addressed with this approach.

We found that TE burden likely has an underdispersed distribution, especially for TEs that are inserted into exons of highly conserved genes and expected to exert large fitness impacts. On the other hand, intergenic TEs near essential genes also have an underdispersed distribution. This observation could be driven by deleterious mechanisms of TEs that could impair host fitness from a distance to genes, such as insertion into regulatory elements or through the spreading of repressive epigenetic marks (reviewed in (Kelleher *et al.* 2020; Choi and Lee 2020)). By comparing various attributes of TE families with and without underdispersed distributions, we found that underdispersing TE families show stronger epigenetic effects and are targeted by more piRNAs generated from the ping-pong cycle. These observations are consistent with the hypothesis that synergism may arise through the deleterious epigenetic effects of TEs. Overall, our discoveries empirically supported the theoretically predicted synergistic fitness effects of TE insertions.

In addition to purifying selection with synergistic epistasis, repulsion LD could arise through selective interference among variants that are separated by small genetic distances (Hill and Robertson 1966; Felsenstein 1974; Garcia and Lohmueller 2020). To address this possibility, we estimated LD between pairs of TEs that are of different physical distance on the same chromosome or on different chromosomes (see Materials and Methods). Contrary to the prediction of selective interference, we observed negative LD mainly among TE pairs that are at least 1kb apart (**Figure S7**). In fact, for categories of TEs that we observed to have underdispersed distributions (nonLTR TEs, TEs in exons, and TEs near essential genes), repulsion LD mainly arises among pairs of TEs that are physically separated by at least 0.1Mb (**Figure S7**). Because our analysis excluded TEs that are in or near pericentromeric heterochromatin, where recombination is strongly suppressed, observed negative linkage among TEs that are physically distant are less likely driven by selective interference than by synergistic fitness effects of TEs.

Different from the proposed source of synergistic epistasis of simple mutations (de Visser *et al.* 2011), synergistic fitness effects of TEs have been predicted to arise through unique mechanisms by which TEs impair host fitness. The illegitimate recombination between nonallelic TEs is predicted to lead to an accelerated removal of TEs with increased TE copy number, or synergistic epistasis (Montgomery *et al.* 1987; Langley *et al.* 1988). Under this model, TEs that are prone involved in ectopic recombination should be more likely to exhibit synergistic epistasis (Langley *et al.* 1988; Dolgin and Charlesworth 2008). While we did not find evidence supporting that TEs in genomic regions with high rates of meiotic recombination are more likely to underdisperse, several assumptions of our analysis could have confounded the results. Recombination landscapes vary between individuals (Dumont *et al.* 2009; Comeron *et al.* 2012; Hunter *et al.* 2016) and populations (Samuk *et al.* 2020), and could have been different between our studied Zambian population and the cosmopolitan population from which the recombination rate was estimated (Comeron *et al.* 2012). We also assumed that the rate of ectopic recombination closely mirrors that of homologous recombination (Lichten *et al.* 1987). This assumption has been questioned by the observed lack of TEs at the tip of the *D. melanogaster* X chromosome, where homologous recombination is strongly suppressed and TEs are expected to accumulate (Langley *et al.* 1988; Charlesworth and Lapid 1989).

We also found that TE families with underdispersed distribution do not follow the predictions of the ectopic recombination model, including being larger in abundance (Montgomery *et al.* 1987; Langley *et al.* 1988), longer in length (Petrov *et al.* 2003) or having higher sequence homology within families (Lee and Langley 2010; Petrov *et al.* 2011). These observations could result from our estimated properties of TE families (from the reference genome) are not representative of the studied population (an African population). On the other hand, our results may suggest that predictions of the ectopic recombination model need to be revised by incorporating additional biological details. It is recently proposed that the dependency of ectopic recombination on TE copy number should plateaus when the number of TE insertions in the genome is large, and the process is unlikely limited by the number of potential recombining targets (Kelleher *et al.* 2020). According to this revised model, synergistic epistasis would only arise when TE copy number is below a certain threshold. In addition, the efficiency of recombination is observed to jointly depend on the length and sequence similarities of (reviewed in (Radman and Wagner 1993; Waldman 2008)), as well as the spatial distance between and orientation of, recombing partners (reviewed in (Renkawitz *et al.* 2014)). A model that incorporates these biological details may provide better predictions for the conditions by which synergistic epistasis may arise via ectopic recombination.

TE-mediated spreading of silencing marks is another mechanism from which synergistic epistasis was predicted to arise (Lee and Langley 2010; Lee 2015). In *Drosophila,* this process is initiated by piRNA-directed epigenetic silencing of TEs (Sienski *et al.* 2012; Le Thomas *et al.* 2013) (reviewed in (Czech *et al.* 2018)). Accordingly, many predictions of the model depend on how piRNAs are generated and target TE sequences.

These include predictions that are shared with the ectopic recombination model but not supported by our observations—TE families that are abundant (Lee and Langley 2010; Lee 2015; Lee and Karpen 2017), long (Lee 2015), and homogenous in sequences (Lee and Langley 2010) are more likely to exhibit synergistic epistasis. Again, the complexities of piRNA generation and targeting that are not considered in the current epigenetic effect model could have led to these discrepancies between predictions and observations. For instance, truncated TEs that lost the ability to transcribe would not contribute to piRNA generations through ping-pong cycle (Sienski *et al.* 2012; Olovnikov *et al.* 2013; Shpiz *et al.* 2014). Also, the targeting of TEs by piRNAs is abolished when the sequence divergence between the two is too large (Post *et al.* 2014; Kotov *et al.* 2019) and is particularly sensitive to mismatches at specific positions within the piRNA sequences (Wang *et al.* 2014; Mohn *et al.* 2015). Simple monotonic relationships could not fully capture how the abundance, length, and sequence homology of TE families influence the occurrence piRNA-targeting of TEs and the associated deleterious epigenetic effects.

On the other hand, the epigenetic effect model uniquely predicts that TE families resulting in more extensive spreading of repressive epigenetic marks, which is initiated by the piRNA-mediated silencing of TEs, should more likely to exhibit synergistic fitness effects. Supporting this prediction, we found that TE families with underdispersed distributions have a higher tendency to elicit epigenetic effects. Interestingly, we also found that piRNAs that target TE families with underdispersed distributions are more likely generated through the ping-pong cycle, which is a predicted source from which synergistic fitness effects of TEs arise (Lee and Langley 2010; Choi and Lee 2020). These observations suggest that TE-mediated epigenetic effects likely contribute to the synergistic epistatic interactions among TE insertions, which could drive the stable containment of TE copy number.

It is worth noting that the statistical power for some of our current analyses may be limited due to challenges associated with studying TEs. TEs have a frequency spectrum that is highly skewed towards rare variants (Figure S1, also see (Stewart *et al.* 2011; Nellåker *et al.* 2012; Cridland *et al.* 2013; Kofler *et al.* 2015; Quadrana *et al.* 2016; Laricchia *et al.* 2017)). This low allele frequency would limit the range of possible LD estimates (Sved and Hill 2018), potentially restricting our ability to detect repulsion LD even if synergistic epistasis among deleterious TEs is present. Also, our ability to identify TEs and infer their biological properties (e.g., length and sequence identity) is limited with short-read sequencing data. Some of these limitations may be alleviated in the near future with the growing number of genomes that are sequenced by 3^rd^-generation long reads, which could significantly improve the identification of TEs and the assembly of their sequences (e.g., Debladis *et al.* 2017; Chakraborty *et al.* 2019; Ellison and Cao 2020).

By leveraging population genetic signals to circumvent direct measurements of individual fitness, we provided empirical evidence for the presence of synergistic epistasis among potentially deleterious TE insertions. While we identified that TE-mediated epigenetic effects may result in synergistic epistasis, our observations also suggest a need to incorporate additional biological details to refine predictions for how synergistic fitness effects of TEs may arise. With revised models and the expanding capacity of TE identifications with long-read sequencing, our analysis framework could provide a path forward to investigate the mode, prevalence, and importance of epistatic interactions in the evolutionary dynamics of TEs.

## Acknowledgment

I would like to thank Charles Langley for inspiring this study and Sohail Mashaal for helpful discussions. Andrea Betancourt, Jae Choi, Brandon Gaut, Yuheng Huang, and Kevin Thoronton provided helpful comments on the manuscript. I also appreciate Andrea Betancourt’s careful editing of the manuscript. This work was supported by NIH R00 GM121868.

