## Supplemental Figures for "Synergistic epistasis of the deleterious effects of transposable elements"

**Figure S1.** $\sigma^2/V_a$ of randomly sampled synonymous variants with and without random masking of alleles as missing data

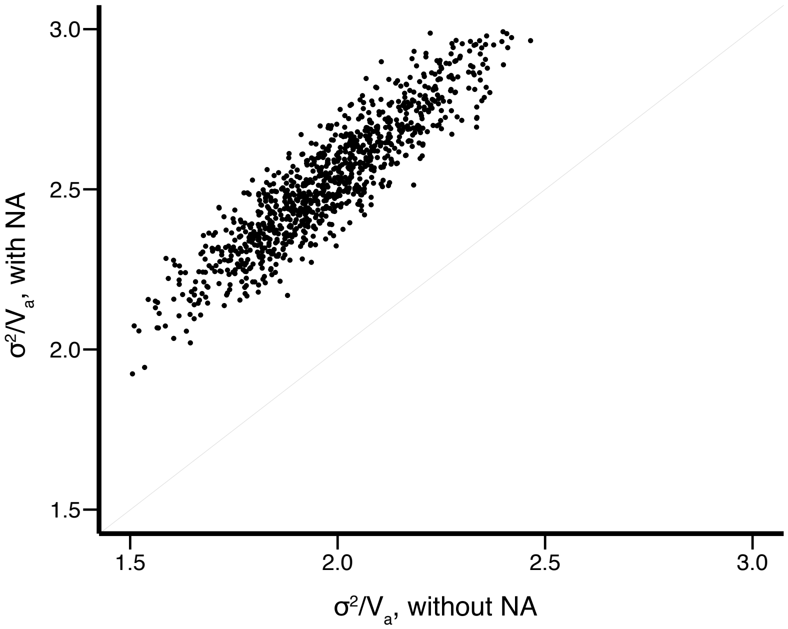

**Figure S2.** The distributions of TE burden for DNA, RNA, TIR, LTR, and nonLTR TEs. Poisson distribution with identical mean is shown as black dots and line.

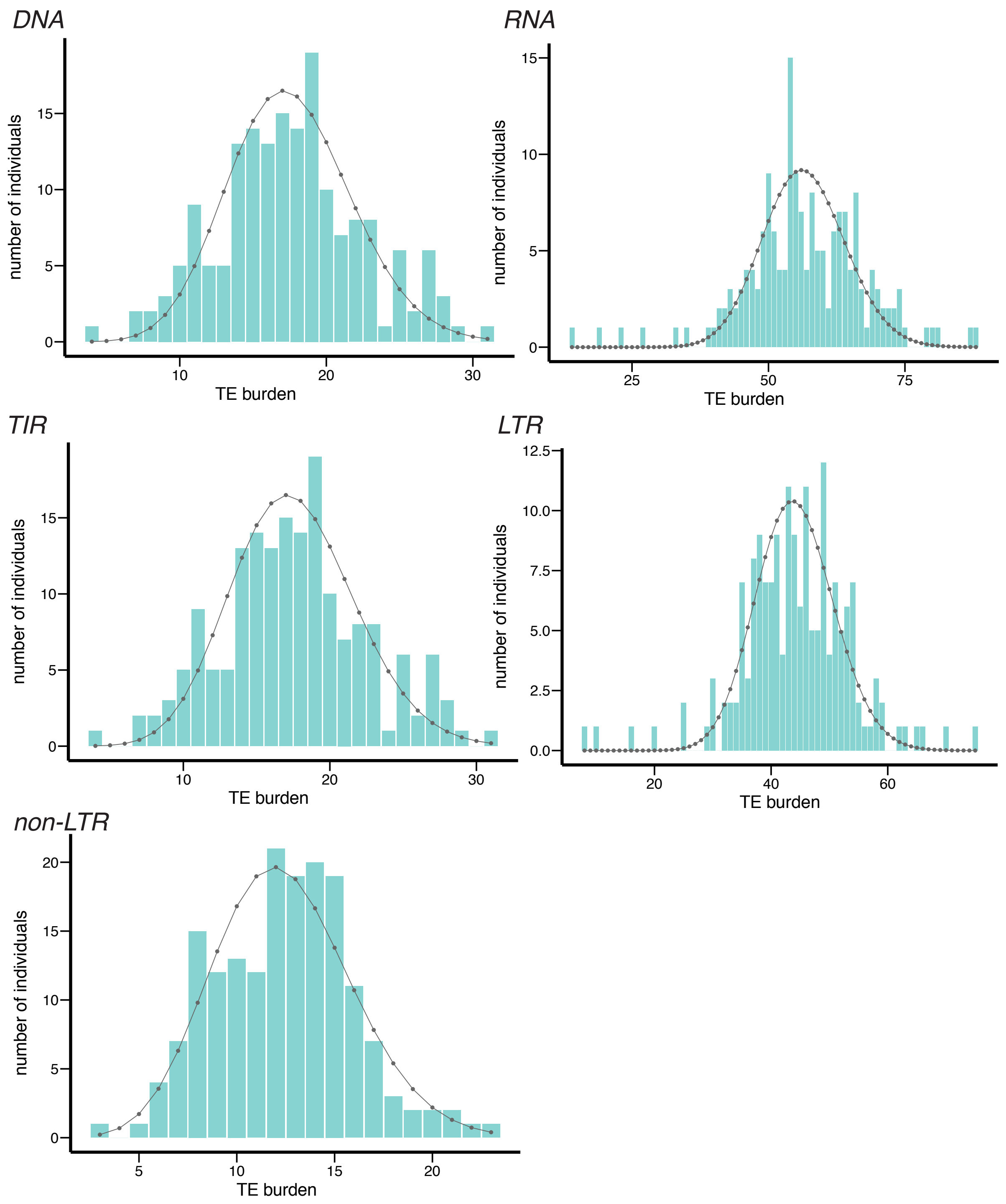

**Figure S3.** The distributions of TE burden for TEs in coding exon, UTR, exon, intron, and intergenic sequences. Poisson distribution with identical mean is shown as black dots and line.

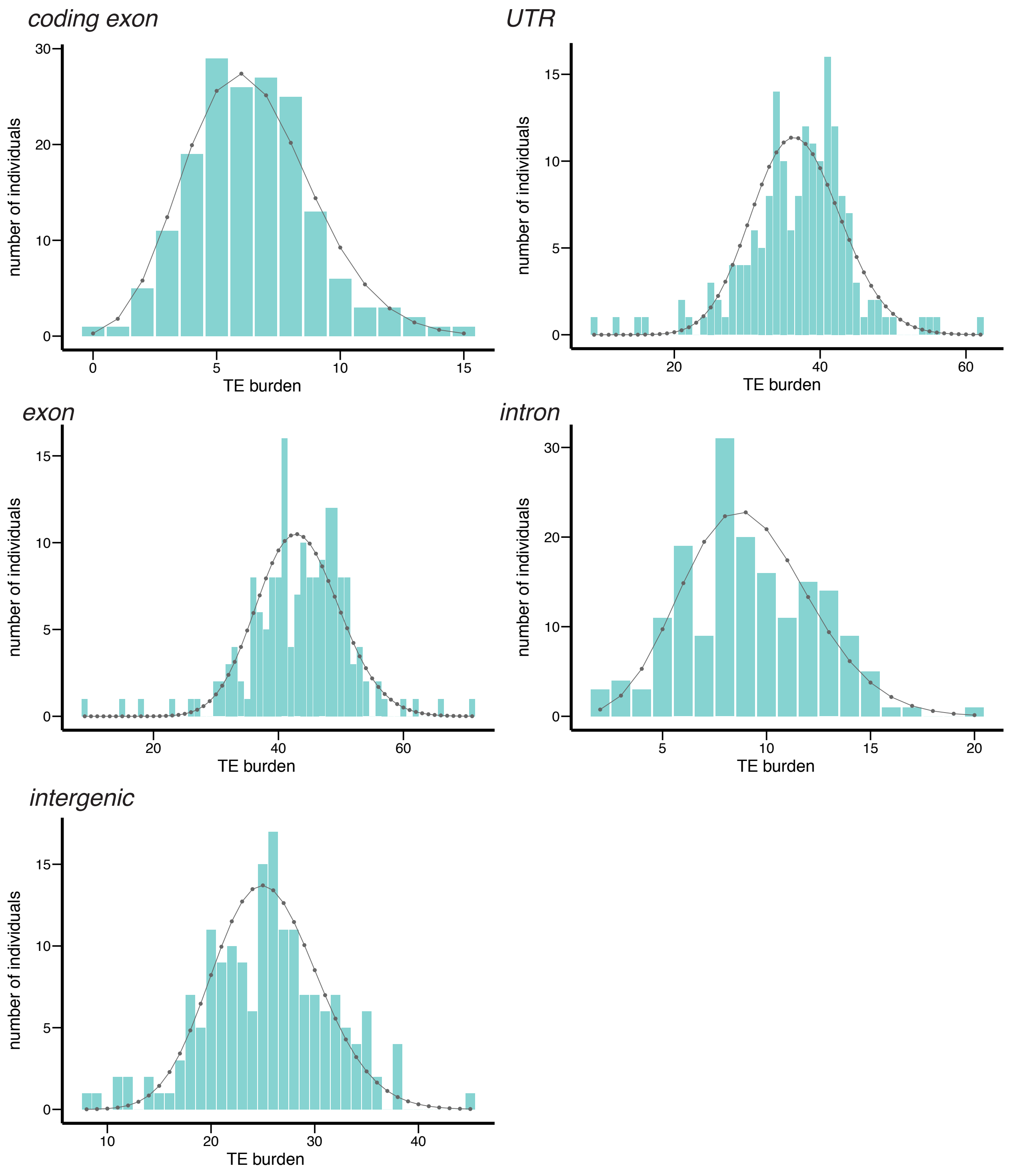

**Figure S4.** The distributions of TE burden for TEs whose nearest genes are under different evolutionary constraints, or *dN/dS* ratio. Poisson distribution with identical mean is shown as black dots and line.

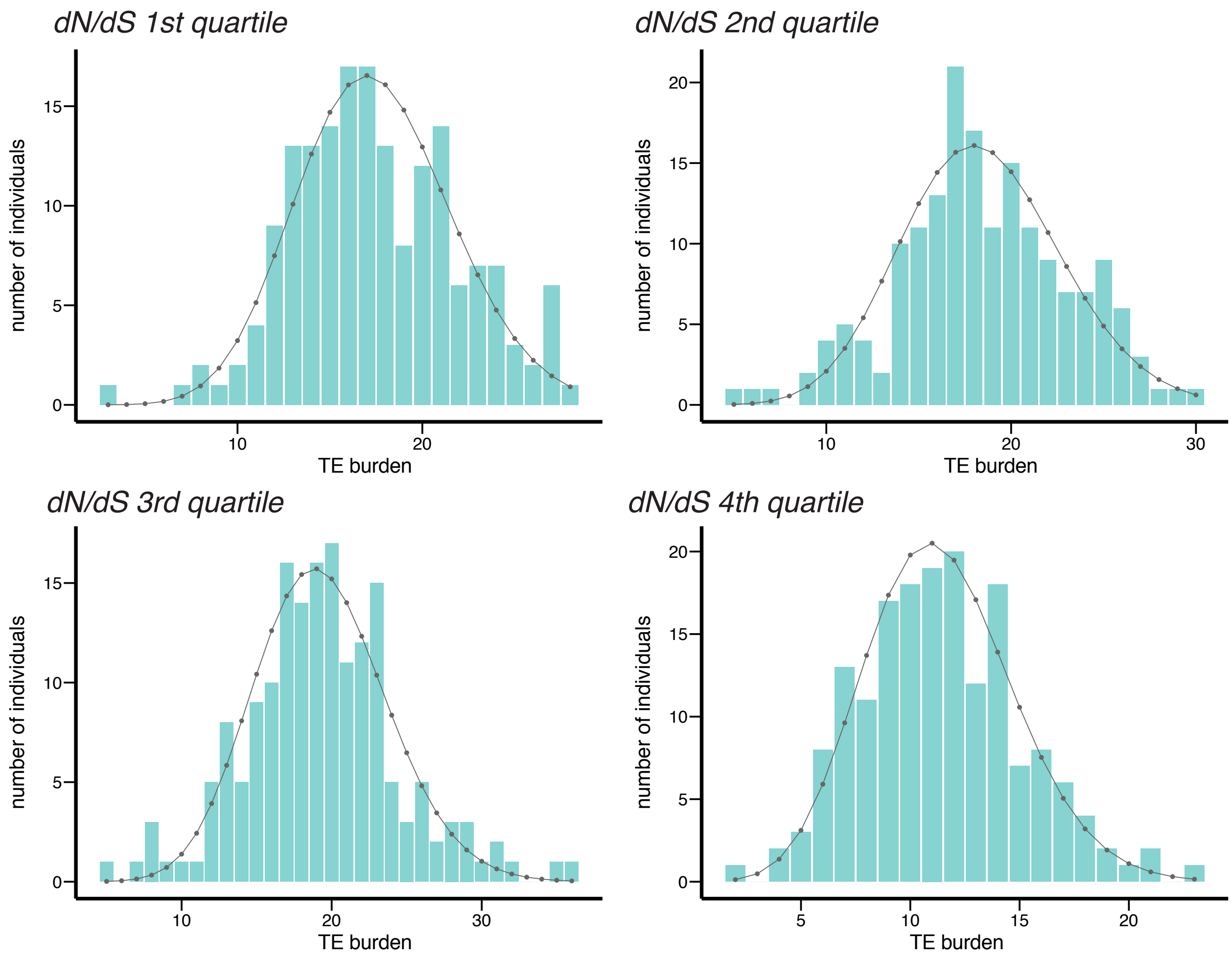

**Figure S5.** The distributions of TE burden for TEs whose nearest genes have different mutant phenotypes, which indicate their essentiality. Poisson distribution with identical mean is shown as black dots and line.

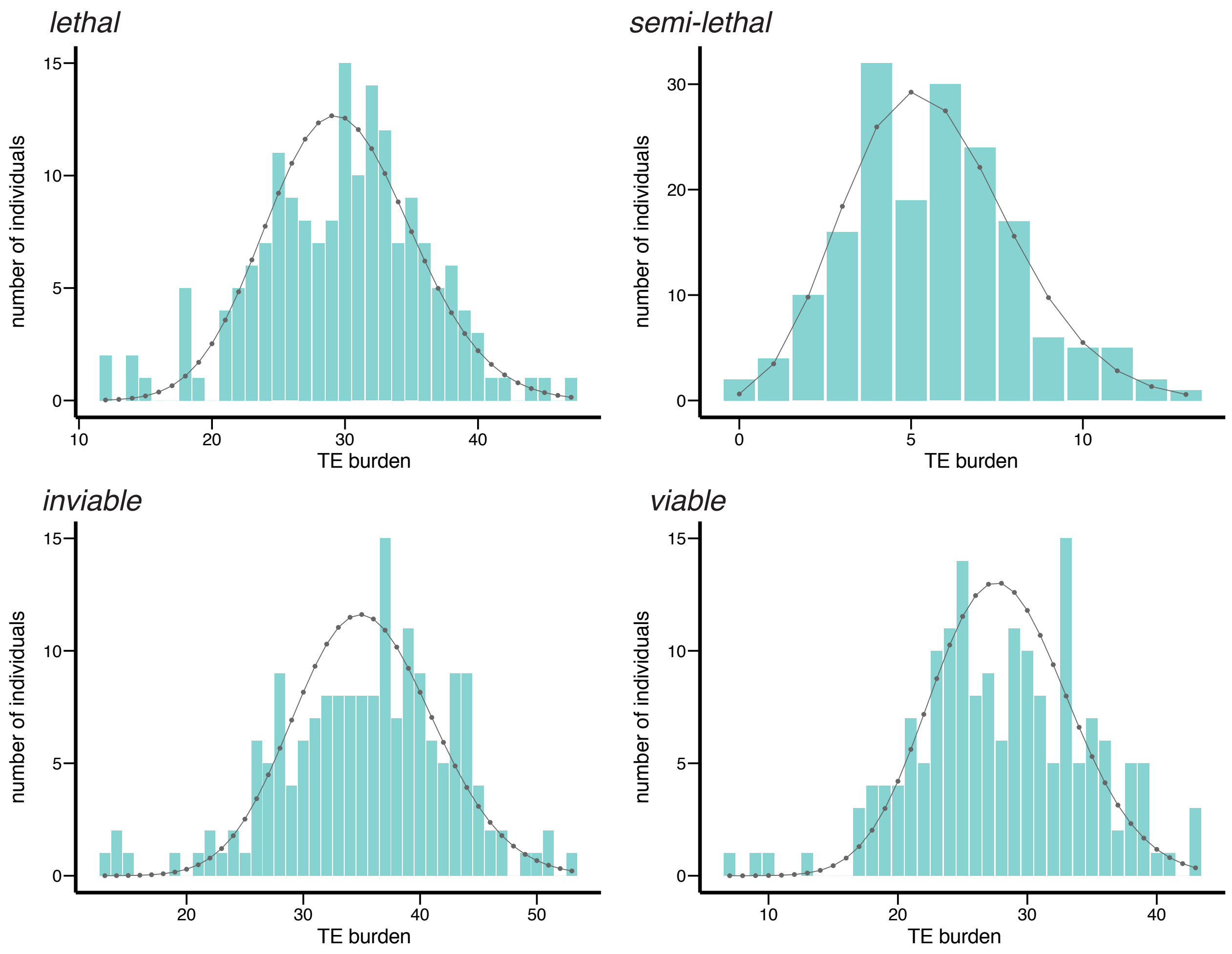

**Figure S6.** The distributions of TE burden for categories of TEs or TE families that were identified to have underdispersed distributions. Poisson distribution with identical mean is shown as black dots and line.

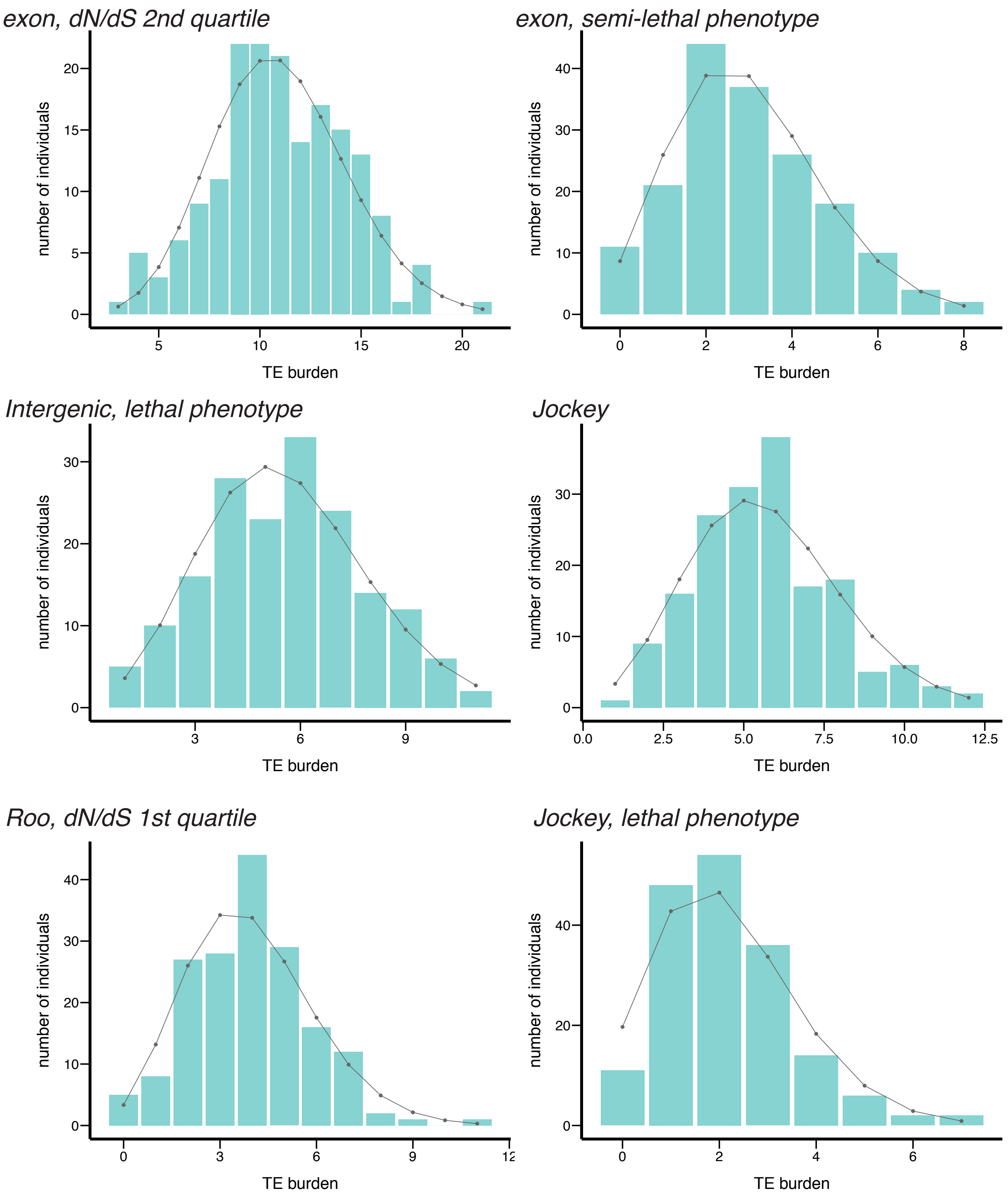

**Figure S7.** Net LD of pairs of TEs that are on the same chromosome of different physical distance or on different chromosomes. Positive LD is shown in gray while negative LD is shown in pink. Error bars show standard errors.

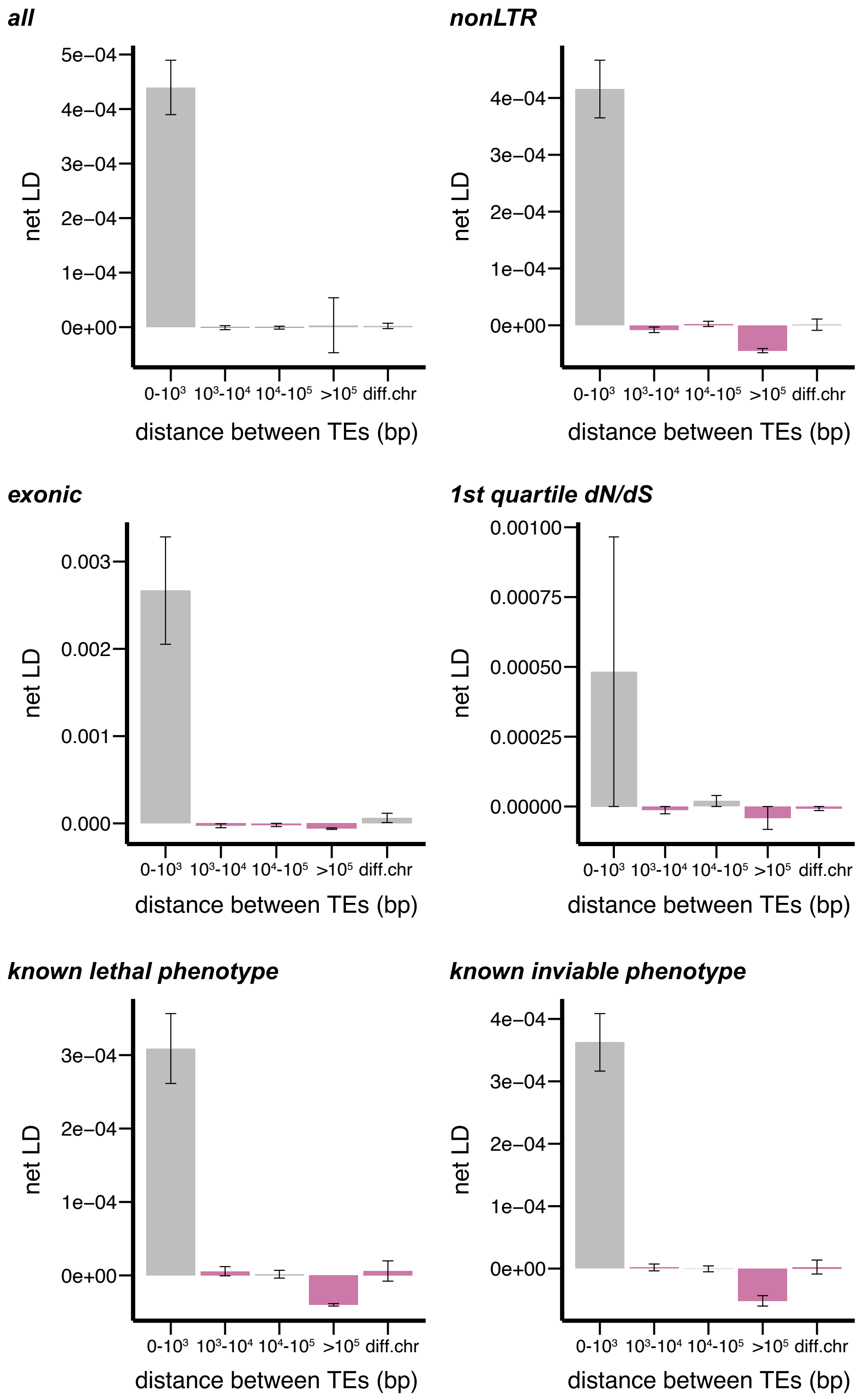
